# Alpha/beta power decreases track the fidelity of stimulus-specific information

**DOI:** 10.1101/633107

**Authors:** Benjamin J. Griffiths, Stephen D. Mayhew, Karen J. Mullinger, João Jorge, Ian Charest, Maria Wimber, Simon Hanslmayr

## Abstract

Massed synchronised neuronal firing is detrimental to information processing. When networks of task-irrelevant neurons fire in unison, they mask the signal generated by task-critical neurons. On a macroscopic level, mass synchronisation of these neurons can contribute to the ubiquitous alpha/beta (8-30Hz) oscillations. Reductions in the amplitude of these oscillations, therefore, may reflect a boost in the processing of high-fidelity information within the cortex. Here, we test this hypothesis. Twenty-one participants completed an associative memory task while undergoing simultaneous EEG-fMRI recordings. Using representational similarity analysis, we quantified the amount of stimulus-specific information represented within the BOLD signal on every trial. When correlating this metric with concurrently-recorded alpha/beta power, we found a significant negative correlation which indicated that as alpha/beta power decreased, our metric of stimulus-specific information increased. This effect generalised across cognitive tasks, as the negative relationship could be observed during visual perception and episodic memory retrieval. Further analysis revealed that this effect could be better explained by alpha/beta power decreases providing favourable conditions for information processing, rather than directly representing stimulus-specific information. Together, these results indicate that alpha/beta power decreases parametrically track the fidelity of both externally-presented and internally-generated stimulus-specific information represented within the cortex.

## Introduction

Neuronal activity fluctuates rhythmically over time. Often referred to as “neural oscillations”, these rhythmic fluctuations can be observed throughout the brain at frequencies ranging from 0.05Hz to 500Hz^1^. When recording from the human scalp, it is the alpha and beta frequencies (8-12Hz; 13-30Hz) that dominate. Alpha/beta activity displays an intimate link to behaviour; engaging in a cognitive task produces a large reduction in the alpha/beta power (amplitude squared). These task-induced power decreases are ubiquitous, and can be observed across species (including humans^2^, macaques^3^, rodents^4^ and cats^5^), sensory modalities (including visual^2^, auditory^6^, and somatosensory^7^ domains), and cognitive tasks (including perception^2,6,7^, memory formation/retrieval^8–10^, and language processing^11^). Given their ubiquity, it stands to reason that these decreases reflect a highly general brain process. While numerous domain-general processes have already been ascribed to alpha/beta oscillations (e.g. idling^12^; inhibition^13,14^), we provide empirical evidence in support of a new perspective: alpha/beta power decreases are a proxy for information processing.

To successfully process information about a stimulus, the brain must be capable of elevating the signal of said stimulus above the noise generated by ongoing neuronal activity^15^. In situations where the ongoing spiking of a large population of neurons is correlated, this is problematic^16^. Mass synchronised spiking generates noise that conceals the comparatively small neuronal signal evoked by the stimulus (see figure 1a), rendering momentary changes in sensory input undetectable^17^ and responses to temporally-extended changes unreliable^18^. Reducing these neuronal “noise correlations”, therefore, can boost the signal-to-noise ratio of an evoked neuronal response to a stimulus. Indeed, numerous studies have demonstrated that the decorrelation of task-irrelevant neuronal firing accompanies engagement in cognitive tasks^18–21^. Given that these noise correlations show a strong positive correlation with the local field potential (LFP)^22^, one may speculate that task-related reductions in alpha/beta LFP^e.g.3^ are (to some degree) a marker of the reduction of noise correlations. Such a hypothesis would explain why reductions in alpha/beta power are associated with the successful execution of a wide range of cognitive tasks, from visual perception^2^ to memory retrieval^23^.

**Figure 1.**
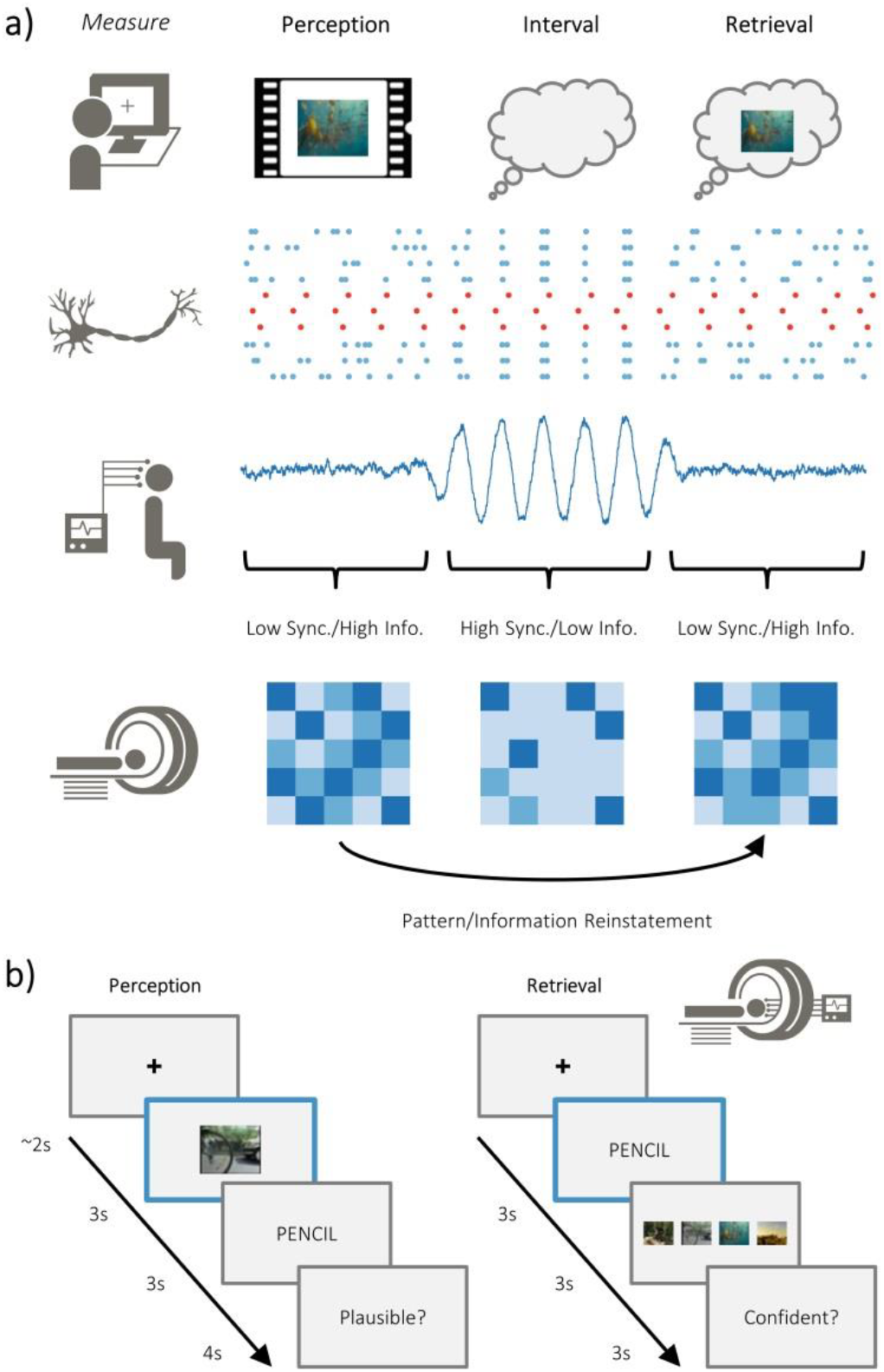
Overview of hypotheses and paradigm. a) The brain is capable of representing stimulus-specific information through neural patterns that are consistent regardless of whether the stimulus is externally or internally generated (i.e. perceived or retrieved; top). On a neuronal level, populations that code for the stimulus (in red) need to generate signal greater than ongoing neuronal noise (in blue). When the neuronal noise correlates (i.e. arises at the same time; during the ‘interval’), the signal-to-noise ratio is reduced and stimulus specific information is limited. These noise correlations may be reflected in macroscopic measures of electrophysiological activity, where periods of highly synchronised firing is accompanied by periods of high amplitude activity. Under this assumption, high amplitude activity would reflect an attenuation of the processing of stimulus-specific information. Stimulus-specific information can be measured using fMRI to look at pattern similarity during perception and pattern reinstatement during memory retrieval. b) Participants completed an associative memory task while undergoing simultaneous EEG-fMRI recordings. Participants were asked to vividly associate a video with a word, and then rate how plausible (i.e. believable) the imagined association was. Later, they were cued with the word and tasked with recalling the associated video. After selecting the associated video, they were asked to judge how confident they felt about their decision.

Here, we test the hypothesis that alpha/beta power decreases are a proxy for information processing^24^. Specifically, we predict that as the amount of stimulus-specific information within the cortex increases, concurrently-recorded measures of alpha/beta power will decrease. Twenty-one participants took part in an associative memory task whilst simultaneous EEG-fMRI recordings were obtained (see figure 1b). On each trial, participants were presented with one of four videos, followed by a noun, and asked to pair the two. Later, participants were presented with the noun and asked to recall the associated video (which would lead to the reinstatement of stimulus-specific information about the video^25^). We first conducted representational similarity analysis (RSA) on the acquired fMRI data to quantify the relative distance between neural patterns of matching and differing videos. This provides a data-driven and objective measure of stimulus-specific information present during a single trial. We then derived alpha/beta power from the concurrently recorded EEG and correlated the observed power with our measure of stimulus-specific information on a trial-by-trial basis. Foreshadowing the results reported below, we found that alpha/beta power decreases negatively correlated with the amount of stimulus-specific information. Importantly, we find evidence for this during both the perception and retrieval of these videos, providing a conceptual replication of our results and supporting the domain-general nature of our hypothesis.

## Results

### Detecting stimulus-specific information in BOLD patterns

Our first step was to derive a measure of stimulus-specific information from the acquired fMRI data. To this end, we used searchlight-based representational similarity analysis (RSA) to quantify the overlap in BOLD patterns for matching videos, and contrasted this against the overlap between differing videos. We interpret the difference in overlap between matching and differing videos as the amount of stimulus-specific information present on a single trial, under the assumption that any similarity that can only be explained by matching stimuli represents information specific to that stimulus. To evaluate whether the quantity of stimulus-specific information was meaningful within a searchlight, the observed measure of information was contrasted against the null hypothesis (i.e. that BOLD pattern overlap for matching videos is the same as BOLD pattern overlap for differing videos) in a one-sample, group-level t-test.

For the perceptual task, stimulus-specific information was quantified by computing the representational distance between every pair of perceptual trials. Searchlight analysis revealed a significant increase in stimulus-specific information relative to chance bilaterally in the occipital lobe (p_FWE_ < 0.001, k = 9911, peak MNI: [x = - 30, y = −57, z = −2], Cohen’s d_z_ = 1.79) [see figure 2a-b]. A frontal central cluster (p_FWE_ < 0.001, k = 113, peak MNI: [x = 12, y = −16, z = 50], Cohen’s d_z_ = 0.28) and a left temporal cluster (p_FWE_ = 0.003, k = 64, peak MNI: [x = −48, y = −1, z = 18], Cohen’s d_z_ = 0.81) also demonstrated a significant increase in stimulus-specific information. These results demonstrate that stimulus-specific information is represented within the cortex during visual perception, and provide a region of interest that yields a meaningful measure of stimulus-specific information for our central analysis.

**Figure 2.**
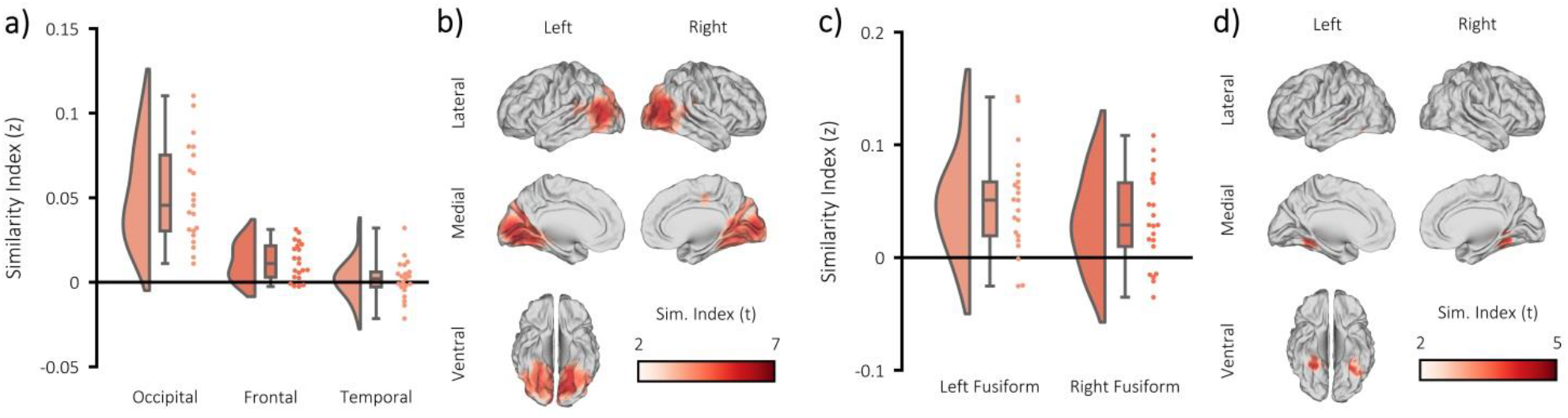
fMRI RSA searchlight analysis. a) raincloud plot depicting the degree to which matching and differing stimuli could be distinguished from one another during visual perception, per participant (single dots), within the three significant clusters (p < 0.05). b) brain map depicting the cluster where matching and differing stimuli could be distinguished from one another during visual perception. c) raincloud plot depicting the degree to which matching and differing stimuli could be distinguished from one another between encoding and retrieval, per participant (single dots), within the two significant clusters (p < 0.001). d) brain map depicting the cluster where matching and differing stimuli could be distinguished from one another during visual memory retrieval.

For the retrieval task, stimulus-specific information was quantified by comparing every retrieval pattern with every perceptual pattern. This approach is sensitive to the reinstatement of veridical information about a successfully recalled stimulus^25^. As we would not anticipate that any stimulus-specific information is present in the BOLD signal when the correct stimulus is not recalled, this analysis was restricted to trials where the paired associate was successfully recalled. Searchlight analysis revealed a significant increase in reinstated stimulus-specific information relative to chance in the right fusiform gyrus (p_FWE_ < 0.001, k = 313, peak MNI: [x = 30, y = - 46, z = −14], Cohen’s d_z_ = 1.07) and left fusiform gyrus (p_FWE_ < 0.001, k = 456, peak MNI: [x = −45, y = −37, z = −6], Cohen’s d_z_ = 0.69) [see figure 2c-d]. These results demonstrate that stimulus-specific information is reinstated during the retrieval of a video, and provide a region of interest that yields a meaningful measure of stimulus-specific information for the central analysis of the memory task.

### Alpha/beta power decreases accompany task engagement

We then measured the degree to which alpha/beta power drops during task engagement. As such an effect is perhaps the most ubiquitous effect in studies of task-related scalp EEG activity, it provides a strong benchmark for the quality of our EEG data (which has the potential for distortion by MRI-related artifacts^26^). For both the perceptual and retrieval trials, the time-series of every source-reconstructed virtual EEG electrode was decomposed into alpha/beta power using 6-cycle Morlet wavelets and baseline-corrected using z-transformation. In the first instance, post-stimulus power (500 to 1500ms) was contrasted against pre-stimulus power (−1000 to 0ms) in a cluster-based, permutation t-test (for perceptual and retrieval trials separately). Unsurprisingly, we found a significant decrease in alpha/beta power following stimulus presentation in both the perceptual (p < 0.001, Cohen’s d_z_ = 0.99; see figure 3a-c) and retrieval (p < 0.001, Cohen’s d_z_ = 0.87; not visualised) tasks. No region exhibited an increase in alpha/beta power during this window.

**Figure 3.**
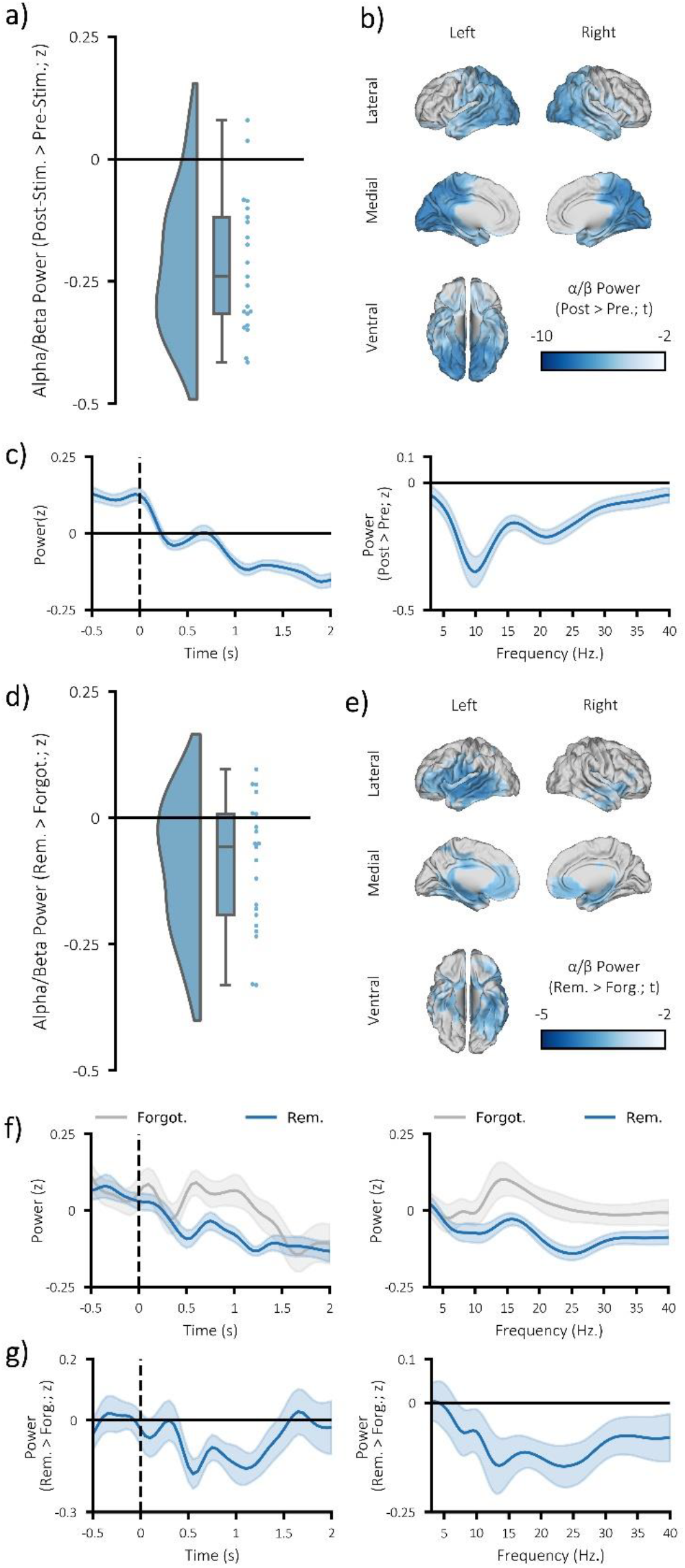
Alpha/beta power decreases during visual perception and memory retrieval. a) Raincloud plot displaying the difference in pre-(−1000ms-0ms) and post-stimulus alpha/beta power (500-1500ms; 8-30Hz) during visual perception, each dot represents a single participant (p < 0.001). b) brain map of the post-stimulus > pre-stimulus difference in alpha/beta power during visual perception (non-significant virtual electrodes are masked). c) the time course of the mean alpha/beta power during perception (left; across participants; 8-30Hz; shaded error bar represents standard error of the mean) and difference in frequency spectrum for post-stimulus > pre-stimulus power during perception (right; across participants; shaded error bar represents standard error of the mean; averaged across all virtual electrodes within significant cluster). d) Raincloud plot displaying the difference post-stimulus alpha/beta power (500-1500ms; 8-30Hz) for successfully recalled pairs relative to forgotten pairs, each dot represents a single subject (p < 0.05). e) brain map of the memory-related difference in alpha/beta power (non-significant virtual electrodes are masked). f) the time course of the mean alpha/beta power (left; across participants; 8-30Hz; shaded error bar represents standard error of the mean) and frequency spectrum (right; across participants; shaded error bar represents standard error of the mean) for remembered and forgotten (in blue and grey respectively; averaged across all virtual electrodes within significant cluster). g) the time course of the mean difference in alpha/beta power (left; across participants; 8-30Hz; shaded error bar represents standard error of the mean) and difference in frequency spectrum (right; across participants; shaded error bar represents standard error of the mean) for remembered relative to forgotten pairs (averaged across all virtual electrodes within significant cluster).

We then asked whether alpha/beta power decreases are not only predictive of task engagement, but also task success. In other words, is the reduction in alpha/beta power greater when memories are successfully recalled? As in the above paragraph, this is not a novel idea and has been demonstrated many times prior^23^. Nevertheless, we wanted to further demonstrate the robustness of our acquired EEG data. To this end, the post-stimulus alpha/beta power (500-1500ms; 8-30Hz; matching previously-reported windows of retrieval-related memory effect^23^) for remembered trials was contrasted with that of forgotten trials in a cluster-based, permutation t-test. Matching earlier reports, we found a significant reduction in alpha/beta power for recalled pairs, relative to forgotten pairs (p = 0.013, Cohen’s d_z_ = 0.57; see figures 3d-g). These power decreases were localised to the late visual ventral stream (including the region within the fusiform gyrus where stimulus-specific information could be identified), as well as other parts of the memory network^27^ (including the medial temporal lobe and medial prefrontal cortex). No region exhibited an increase in alpha/beta power during this window. In sum, these results demonstrate that alpha/beta power decreases accompany the engagement in, and successful execution of, cognitive tasks.

### Alpha/beta power decreases track the fidelity of stimulus-specific information

We then addressed our central question: do alpha/beta power decreases parametrically track the fidelity of stimulus-specific information? For each subject, a single trial measure of stimulus-specific information was computed by comparing the trial pattern within the region of interest (i.e. the significant clusters identified in the fMRI searchlight analysis; see figure 2b) to patterns of matching and differing videos. For the perceptual data, this approach involved computing the representational distance for every pair of perceptual trials. These distances were then correlated with a unique model for each trial that stated representational distance for stimuli matching the stimulus presented would be zero and representational distance for stimuli differing from the stimulus presented on this trial would be one. The resulting correlation coefficient was Fisher z-transformed to provide a normally-distributed metric of stimulus-specific information for each trial. Alpha/beta power within the region that housed stimulus-specific information was calculated and averaged over virtual electrodes, frequency and time. The metric of stimulus-specific information was then correlated with alpha/beta power across trials (see figure 4a). The resulting correlation coefficient was then Fisher z-transformed (to approximate a normal distribution). These Fisher z-values were contrasted against the null hypothesis (there is no correlation; z = 0) across participants in a one-sample t-test. We found a significant negative correlation (p = 0.044, Cohen’s d_z_ = 0.37), where a reduction in alpha/beta power was accompanied by an increase in stimulus-specific information (see figure 4b). When examining which virtual EEG electrodes showed the greatest negative correlation, we found evidence to suggest this effect was primarily localised to the occipital alpha/beta power (p = 0.046, Cohen’s d_z_ = 0.50; see figure 4c), overlapping with the regions where stimulus-specific information was detected (see figure 2b). This result demonstrates that alpha/beta power tracks the fidelity of stimulus-specific information during visual perception.

**Figure 4.**
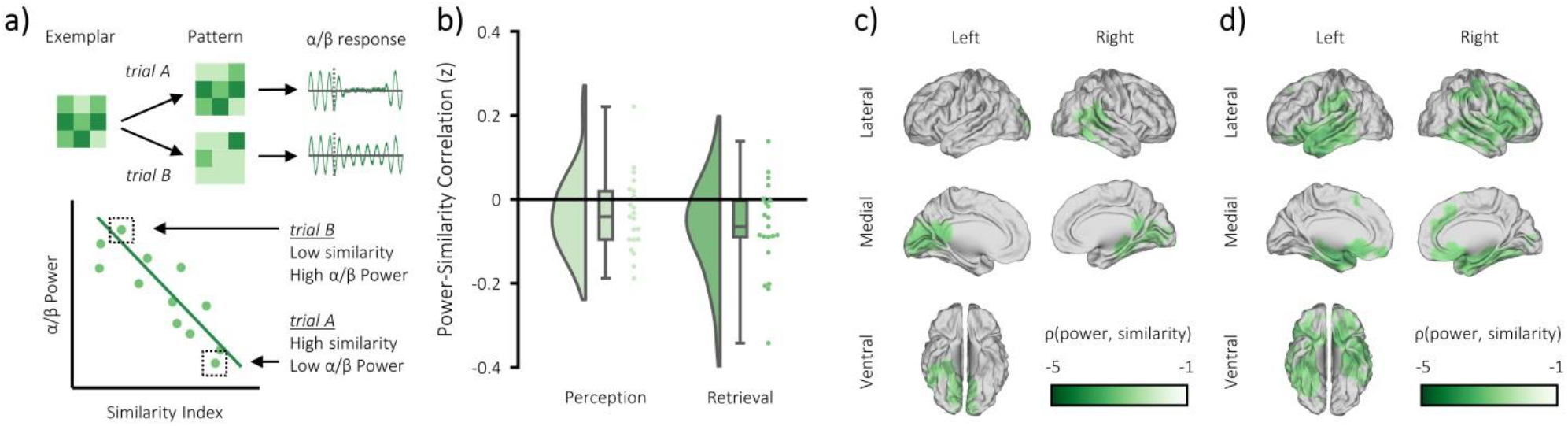
Alpha/beta power decreases track the fidelity of stimulus-specific information. (a) infographic depicting hypotheses and analytical approach. We anticipated that the more a pattern represented matching stimuli relative to differing stimuli, the greater the post-stimulus decrease in alpha/beta power would be. (b) Raincloud plot displaying the correlation between alpha/beta power and stimulus-specific information during visual perception and memory retrieval (each dot represents a single participant; p < 0.05. cf. null hypothesis). (c) brain map of the correlation between alpha/beta power at each virtual electrode with the measure of stimulus-specific information during visual perception (non-significant virtual electrodes are masked (d) brain map of the correlation between alpha/beta power at each virtual electrode with the measure of stimulus-specific information during memory retrieval (non-significant virtual electrodes are masked).

We then aimed to replicate this effect in the retrieval task, working on the assumption that if alpha/beta power decreases are a proxy for information processing, the phenomenon should generalise across cognitive tasks. The correlation analysis was conducted separately for remembered and forgotten pairs to avoid a spurious correlation driven by memory-related differences in the decreases of alpha/beta power and increases of stimulus-specific information for remembered compared to forgotten trials. Representational distance was calculated between each single trial at retrieval and all trials at perception within the region of interest (i.e. the significant clusters identified in the fMRI searchlight analysis; see figure 2d), and then correlated with a model that stated that representational distance for perceived stimuli matching the retrieved stimulus on this trial would be zero and representational distance for perceived stimuli differing from the retrieved stimulus on this trial would be one. The remainder of the analysis is the same as described above. In line with the previous result, we found a significant negative correlation for remembered trials (p = 0.004, Cohen’s d_z_ = 0.61), where a reduction in alpha/beta power was accompanied by an increase in stimulus-specific information (see figure 4b). When examining which virtual EEG electrodes showed the greatest negative correlation, the largest cluster was localised to alpha/beta power in the late visual ventral stream and medial temporal/frontal regions (see figure 4d), overlapping with the regions where memory-related decreases in alpha/beta power was observed (see figure 2d). No effect was observed when conducting this analysis on forgotten trials (p = 0.213, Cohen’s d_z_ = 0.18), perhaps because little stimulus-specific information will be represented when the memory cannot be retrieved. These results support the earlier conclusion that alpha/beta power decreases parametrically track the fidelity of stimulus-specific information.

Notably, alpha/beta power and BOLD signal have previously been shown to correlate negatively^28^ (also see supplementary materials). To rule out the possibility that the link between alpha/beta power and stimulus-specific information is driven simply by a change in BOLD signal, we ran a partial correlation controlling for the confound of BOLD intensity. In line with our earlier findings, we found a significant negative correlation between alpha/beta power and stimulus-specific information during perception (p = 0.028, Cohen’s d_z_ = 0.44) and memory retrieval (p = 0.011, Cohen’s d_z_ = 0.56), indicating that the observed correlation cannot be explained by BOLD intensity alone.

Additionally, alpha/beta power decreases correlate with the participant’s confidence of recalling the video-word association (see supplementary materials). To rule out the possibility that alpha/beta power decreases are a confidence signal that increases as more stimulus-specific information is recalled about an association, we ran a partial correlation probing the link between alpha/beta power and similarity during memory retrieval while controlling for confidence. Matching the previous result, we found a significant negative correlation (p = 0.006, Cohen’s d_z_ = 0.59), indicating that the observed correlation between alpha/beta power decrease and stimulus-specific memory reinstatement cannot be explained by confidence alone.

In sum, these results suggest that alpha/beta power parametrically decreases as the amount of stimulus-specific information represented within the cortex increases. While the chance of uncovering these two central effects were not much smaller than the threshold for significance (α = 0.05; where observed p values were 0.044 and 0.004), it is worth noting that the joint probability of finding both effects in support of our hypothesis was substantially smaller (p < 0.001).

### Alpha/beta power decreases do not represent perceived or retrieved information

Lastly, we asked whether the observed negative correlation between alpha/beta power and stimulus-specific information could be explained by the fact that alpha/beta power, rather than providing favourable conditions for the brain to represent activity, actually represents information itself. To test this hypothesis, we conducted spatiotemporal representational similarity analysis (i.e. across virtual electrodes, time windows [500 to 1500ms, in steps of 100ms] and frequency bins [8 to 30Hz, in steps of 1Hz]) within the regions where stimulus-specific information was identified in the BOLD signal during perception and memory retrieval. By restricting analysis to regions where we had previously detected stimulus-specific information, we maximise our chance of finding an effect. Despite this extremely liberal approach, a cluster-based, permutation t-test found no evidence to suggest that alpha/beta power represents stimulus-specific information during perception (p = 0.548, Cohen’s d_z_ = 0.03) or retrieval (p = 0.579, Cohen’s d_z_ = 0.04). Notably, the frequentist nature of this test means we cannot conclude that alpha/beta power does not represent information, but rather that there is insufficient evidence to conclude that alpha/beta power represents information. To address this limitation, we ran a Bayesian one-sample t-test to probe the nature of the evidence in favour of the null hypothesis. The continued use of the region of interest switches the test from a liberal test of the alternative hypothesis (alpha/beta represents information) to a conservative test of the null hypothesis (alpha/beta does not represent information). Bayesian one-sample t-tests revealed moderate evidence in favour of the null hypothesis for the perceptual (BF_10_ = 0.230) and retrieval tasks (BF_10_ = 0.232). These results suggest that the observed relationship between alpha/beta power decreases and stimulus-specific information cannot be explained by the hypothesis that alpha/beta power itself represents information. Rather these results suggest that alpha/beta power decreases are a marker of the fidelity of stimulus-specific information.

## Discussion

Here, we provide the first empirical evidence that alpha/beta power decreases track the fidelity of stimulus-specific information represented within the cortex. We correlated simultaneously-recorded alpha/beta power (as measured using scalp EEG) with a metric of stimulus-specific information (as quantified using representational similarity analysis [RSA] on fMRI data) on a trial-by-trial level. As stimulus-specific information increased, alpha/beta power decreased, regardless of whether the information was externally presented or internally generated. Further analysis revealed that this effect is not driven by the fact that alpha/beta power decreases represent information, suggesting instead that they provide conditions which are beneficial for information processing.

Our central finding demonstrates that as alpha/beta power decreases, the fidelity of stimulus-specific information within the cortex increases. Task-related decreases in alpha/beta power are observable across tasks^2,6–11^, sensory modalities^2,6,7^, and species^2–5^. Given their ubiquity, it stands to reason that they reflect a highly general cognitive process. While others have attributed similar results to idling^12^ or inhibition^13^, we provide evidence that these alpha/beta power decreases are a proxy for information processing. This supports the idea that a reduction of neuronal noise correlations (which map onto local field potential; LFP^22^) can facilitate the representation of information^16^. Numerous studies have demonstrated that task-irrelevant correlated activity between pairs of neurons is detrimental to stimulus processing^15,29^ – particularly for large networks of correlated neurons^16^ that, incidentally, are more likely to be detected in the LFP. As our conclusion works on the assumption that a reduction in LFP equates to a reduction in noise correlations, we open up an interesting new question: do measures of noise correlations directly map onto an objective and parametric measure of stimulus-specific information? Addressing this question would further strengthen the view that reducing underlying noise can boost the information processing capabilities of the cortex.

Following the hypothesis that alpha/beta power decreases are a proxy for reductions in noise correlations, one would predict that alpha/beta power decreases do not carry representational information about a stimulus. Rather, they provide favourable (i.e. reduced noise) conditions in which another mechanism can allow the internal representation of said stimulus to come forth. In line with this hypothesis, we found moderate evidence to suggest that alpha/beta power decreases do not carry any stimulus-specific information during the perception or retrieval of the visual stimuli. As such, one would view alpha/beta power decreases as a marker for the potential for information processing, rather than representing information.

The central results can also be explained by information theory^30^. Information theory proposes that little information can be gathered from a highly predictable input (e.g. a network of highly correlated, spiking neurons) – if you can predict an upcoming event, you must already know details about the event. In contrast, a lot of information can be gathered from unpredictable inputs (e.g. uncorrelated spiking neurons) – you learn a lot from a completely novel experience. It has been theorised that desynchronisation of alpha and beta oscillations reduces the predictability of neuronal firing and hence boosts information processing abilities^24^. For example, earlier work has demonstrated that tasks which involve greater semantic elaboration (i.e. greater information processing) produce greater alpha/beta power decreases^9^. Our key result fits neatly within this framework as we find that alpha/beta power decreases parametrically increase with the presence of stimulus-specific information. Moreover, our finding that alpha/beta power does not directly represent stimulus-specific information fits with this idea, as these power decreases are theorised to allow complex neuronal patterns to emerge rather than generate the complex patterns themselves. Taken together, one could speculate that alpha/beta power decreases allow for the rich representation of stimulus-specific information by reducing the predictability of neural firing patterns. Notably, the information theoretic interpretation (i.e. predictable firing is bad for information processing) is highly similar to the idea that correlated firing (i.e. noise correlations) is bad for information processing because correlations are inherently predictable. This opens an exciting new line of questions which ask whether metrics of information are benefitted from the indiscriminate attenuation of synchronised neuronal firing (i.e. reducing redundancy) or from the selective attenuation of neurons that contribute to noise correlations (i.e. boosting signal-to-noise).

As alluded to earlier, high-amplitude alpha oscillations have previously been interpreted as a marker for inhibition^12–14^. One may wonder then, how can the current results and theory be reconciled with these established accounts? Quite simply, we view our account and the existing inhibition accounts as two sides of the same coin. Earlier accounts focus upon how alpha power increases reflect inhibition, our framework focuses on the complementary idea that alpha power decreases boost information representation through disinhibited networks. Importantly, we expand on these earlier accounts by demonstrating that alpha/beta power does not simply reflect a binary division between inhibition and disinhibition. Rather, alpha/beta power can parametrically track the degree to which a network can represent information.

Both the causality and directionality of the central result remains open to debate. Perhaps the most critical question is whether alpha/beta power decreases are a prerequisite for information processing. We speculate that this is not the case. Our theoretical interpretation of the results views these power decreases as a means to boost a stimulus’s signal-to-noise ratio by reducing noise correlations. Arguably however, the stimulus’s signal-to-noise ratio can also be boosted by increasing the stimulus’s signal intensity^31^. This would lead us to hypothesise that alpha/beta power decreases are sufficient, though not necessary, for information processing. This hypothesis would explain the size of the per-subject correlation values observed here and in previous studies that linked noise correlations and information processing^29,32^ – if other processes contribute to information processing, the correlation will not be perfect. Indeed, this hypothesis is supported by a study where task-related alpha/beta power decreases were disrupted by transcranial magnetic stimulation^10^ (TMS). In this study, TMS reduced behavioural performance (suggesting that task-related alpha/beta power decreases facilitate information processing), but did not render participants completely incapable of recalling information (suggesting other processes also contribute to information processing). This reasoning generates an interesting question: does brain stimulation impair measures of stimulus-specific information in the BOLD signal by entraining alpha/beta activity? Addressing this question would help to clarify the extent to which alpha/beta activity influences the representation of stimulus-specific information within the cortex.

Our results focused upon visual information processing in humans, which begs the question: does this phenomenon generalise across species and/or sensory modalities? Alpha/beta power decreases are a ubiquitous phenomenon that transcends species^2–5^ and sensory modalities^2,6,7^. Under the assumption that these decreases reflect a common process, we would speculate that alpha/beta power decreases track information processing across species/modalities in the same manner as observed here for visual information in humans. Indeed, previous work has demonstrated that the alpha/beta power decreases that accompany successful retrieval of auditory memories arise in the same region as where temporal patterns of auditory information are decodable^23^, indicating that the principle generalises across modalities. To our knowledge however, no study has attempted to test the idea that information processing in other species is facilitated by alpha/beta power decreases. Perhaps this is an avenue of very fruitful future research.

In this experiment, we focused on the alpha/beta frequencies (8-30Hz) for both theoretical^24^ and pragmatic reasons^26^. This focus does ask however: do the theta and gamma frequencies (3-7Hz; 40-100Hz) relate to information in a similar manner? Both the perception and retrieval of stimuli typically induce power increases in the theta and gamma bands^e.g.33^. These power increases are not overtly congruent with the theories of information processing via neuronal decoupling^15,16,29^ or neuronal unpredictability^24^. As alpha/beta power decreases are proposed to facilitate information processing by reducing noise, however, these theta or gamma power increases could theoretically facilitate information processing through the complementary means of increasing signal strength. For example, the “communication through coherence” hypothesis proposes that neuronal representations of a stimulus are enhanced by an increase in gamma synchronicity^31^. Given that alpha/beta power decreases frequently co-occur with gamma power increases^e.g.34^, one could speculate that these two mechanisms interact such that the former reduces noise while the latter boosts signal to further optimise the efficiency of information processing. As recordings of the theta and gamma frequencies are impaired by the MRI-induced artifacts^26^, a direct test of this theory here is impossible. Nevertheless, future studies can ask whether alpha/beta power decreases interact with other processes or frequencies to enhance the representation of stimulus-specific information in the cortex.

In conclusion, we find evidence to suggest that alpha/beta power decreases track the fidelity of stimulus-specific information represented within the cortex. Given that these alpha/beta power decreases are observed across tasks^2,6–11^, sensory modalities^2,6,7^, and species^2–5^, it stands to reason that they reflect a highly general cognitive process. While earlier theories have linked this phenomenon with idling^12^ and inhibition^13^, our findings suggest they reflect information processing. These power decreases may act as a proxy for information processing either through their link to reduced neuronal noise correlations^15,16,22^ or by reducing the predictability of neuronal activity^24^. These results open numerous avenues for future research, such as how these decreases interact with other neural processes to facilitate the representation of stimulus-specific information, and whether brain stimulation can be used to manipulate the fidelity of information represented within the cortex. Ultimately, these results further illuminate how the ubiquitous phenomenon of task-related alpha/beta power decreases relate to the processing and comprehending of our physical and mental worlds.

## Method

### Participants

Thirty-three participants were recruited. All participants were Native English speakers with normal or corrected-to-normal vision. In return for their participation, they received course credit or financial reimbursement. Twelve of these participants were excluded from analysis: one participant was excluded due to recording issues relating to the MRI scanner, three participants were excluded due to recording issues relating to the EEG system, five participants had insufficient recalled pairs (n<10) following EEG artifact rejection, and three participants had insufficient forgotten pairs (n<10) following EEG artifact rejection. Ethical approval was granted by the Research Ethics Committee at the University of Birmingham, complying with the Declaration of Helsinki.

### Behavioural paradigm

Each participant completed a paired associates task^23,35^ (see fig. 1b). During encoding, participants were presented with a 3 second video or sound, followed by a noun. There was a total of four videos and four sounds, repeated throughout each block. All four videos had a focus on scenery that had a temporal dynamic, while the four sounds were melodies performed on 4 distinct musical instruments. Participants were asked to “vividly associate” a link between every dynamic and verbal stimulus pairing. For each pairing, participants were asked to rate how plausible (1 for very implausible and 4 for very plausible) the association they created was between the two stimuli (the plausibility judgement was used to keep participants on task rather than to yield a meaningful metric, and to ensure that motion in perceptual and retrieval blocks was consistent to aid comparability between tasks). The following trial began immediately after participants provided a judgement. If a judgement was not recorded within 4 seconds, the next trial began. This stopped participants from elaborating further on imagined association they had just created. After encoding, participants completed a 2-minute distractor task which involved making odd/even judgements for random integers ranging from 1 to 99. Feedback was given after every trial. During retrieval, participants were presented with every word that was presented in the earlier encoding stage and, 3 seconds later, asked to identify the associated video/sound from a list of all four videos/sounds shown during the previous encoding block. The order in which the four videos/sounds were presented was randomised across trials to avoid any stimulus-specific preparatory motor signals contaminating the epoch. Following selection, participants were asked to rate how confident they felt about their choice (1 for guess and 4 for certain). Each block consisted solely of video-word pairs or solely of sound-word pairs – there were no multimodal blocks. Each block consisted of 48 pairs, with each dynamic stimulus being presented an equal number of times (i.e. 12 repetitions of each dynamic stimulus). There were 4 blocks in total. After the second block, the structural T1-weight image was acquired, giving participants a chance to rest. Any participant that had fewer than 10 “remembered” or 10 “forgotten” trials after EEG pre-processing were excluded from further analysis. All participants completed the task in the MRI scanner, with fMRI and EEG data acquisition occurring at both encoding and retrieval. Responses were logged using NATA response boxes.

### A note on the auditory variant of paradigm

As mentioned above, all participants completed a variant of the task where the videos were replaced with sounds (as in ^23^) with the aim of replicating the central results in auditory domain. We were unable to dissociate the spatial BOLD patterns of the four melodies, however, meaning we could not derive a measure of stimulus-specific information to test our central hypothesis with. As a result, this aspect of the experiment was discarded.

### Behavioural analysis

Trials were characterised as ‘remembered’ or ‘forgotten’. Remembered trials corresponded to those in which the participant could link the verbal cue to the correct video, and indicated that their decision was not a guess (i.e. confidence rating > 1). Forgotten trials corresponded to those in which the participant could not link the verbal cue to the correct video, or indicated that their decision was a guess (i.e. confidence rating = 1). While earlier studies using this paradigm ^23,35^ have only considered ‘highly confident’ memories (i.e. max confidence rating), we chose a more lax confidence threshold to ensure that sufficient trials of each dynamic stimulus available for the fMRI representational similarity analysis. Under these criteria, participants (on average) correctly recalled 63.4% of the video-word pairs (s.d. 7.5%; range: 47.6-74.5%).

### fMRI acquisition

The magnetic resonance imaging data was acquired using a 3T Philips scanner with a 32-channel SENSE receiver coil at the Birmingham University Imaging Centre (BUIC). Participants were instructed to avoid moving as much as they could, and motion was further restricted by placing foam pads inside the RF coil. Functional volumes consisted of 32 axial slices (4mm thickness) with 3×3mm voxels, providing full head coverage (field of view: 192×192×128mm), acquired through an echo-planar imaging (EPI) pulse sequence (TR=2s, TE=40ms, flip angle of 80°). Four dummy scans were acquired immediately prior to the beginning of each run to allow for magnetic field stabilisation. Eight runs were obtained (4 encoding runs and 4 retrieval runs), each of which acquired 255 volumes plus four dummy scans. A T1-weighted structural image (1×1×1mm voxels; TR = 7.4ms; TE = 3.5ms; flip angle = 7°, field of view = 256 × 256 × 176mm) was acquired after the second block.

### fMRI pre-processing

Pre-processing of the fMRI data was conducted in SPM 12. The functional images first underwent slice time correction, followed by spatial realignment to the first volume of each run. The structural T1-weighted image was then co-registered to the mean image of the functional MRI data. The co-registered T1-weighted image was then segmented. For the univariate analysis (see supplementary materials), the functional and structural images were normalised to MNI space, and then smoothed using a 8×8×8mm full-width at half-maximum (FWHM) Gaussian kernel.

### fMRI representational similarity analysis

Searchlight-based representational similarity analysis (RSA) was conducted using a combination of the MRC CBU RSA toolbox (http://www.mrc-cbu.cam.ac.uk/methods-and-resources/toolboxes/) and custom scripts (https://github.com/benjaminGriffiths/reinstatement_fidelity). Representational distance was quantified as the cross-validated Mahalanobis (CVM) distance^36,37^, which provides an unbiased measure of pattern dissimilarity^37^. The CVM approach takes a training dataset and finds weights that maximises the Euclidean distance between two stimuli. These weights are then applied to a testing dataset, and the weighted Euclidean (i.e. cross-validated Mahalanobis) distance is calculated between stimuli. For the analysis of the perceptual task, the time-corrected and spatially-realigned fMRI data was demeaned and then split into two partitions, with the first partition containing data from the first block and the second partition containing data from the second block. The first partition served as training data for calculating CVM distance on the second partition, and the second partition served as training data for calculating distance on the first partition. CVM distance was computed between every stimulus pattern at encoding. The derived CVM distance was then correlated with a hypothesised model, which stated that (i) there would be a perfect correlation (r = 1) between the representation of each repetition of the same video, and (ii) there would be no correlation (r = 0) between the representation of differing videos. Spearman’s correlation was used based on the ordinal nature of the hypothesised model. The resulting correlation co-efficient was then corrected using the Fisher z-transform to approximate a normal distribution. This analysis was conducted across the whole brain using searchlights with a radius of 10mm (i.e. 121 voxels). Searchlights that contained less than 60% of these 121 voxels (e.g. searchlights in the most lateral areas of the neocortex) were discarded from analysis. The Fisher z-value of each searchlight was placed in a brain map, at the centre voxel of the searchlight. For statistical inference, the resulting brain maps of each subject were analysed in a second-level one-sample t-test. The resulting group-level brain map was thresholded in SPM using p_uncorr._ < 0.001 and a cluster extent of k = 10.

For the retrieval task, this analysis was adapted slightly. The cross-validation method used above assumes that each representation of the same video is identical, and while this is true for perception (participants always viewed one of the four identical video clips), the same is not true for retrieval (each memory consists of a unique word-video pair). To address this concern, trials that contained the same video were averaged together to maximise the video-stimulus “signal” and minimise the word-stimulus “noise”. These mean patterns were then subjected to the same analysis as above. Weights maximising the Euclidean distance between each mean pattern were calculated on a training dataset, and applied to the testing dataset to allow the calculation of the CVM distance. This was conducted between every pattern at both perception and retrieval. The observed distances were then correlated with a hypothesised model, which stated that (i) there would be a perfect correlation (r = 1) between the mean representation of a video at retrieval and the mean representation of the same video at perception, and (ii) there would be no correlation (r = 0) between the mean representation of a video at retrieval and the mean representations of differing videos at perception. Any cases of perception-perception or retrieval-retrieval similarity were excluded from this model, meaning this model isolates the effects of memory reinstatement. The approaches to searchlight analysis and statistical inference were identical to those described in the previous paragraph.

### EEG acquisition

The EEG was recorded using a MR compatible Brain Products system (Brain Products, Munich, Germany) and a 64-electrode cap with a custom layout (including an EOG and ECG channel). As movement within the scanner has been shown to profoundly impair EEG data quality ^26^, motion sensors were attached to the EEG cap to assist in the attenuation of movement-related EEG artifacts ^38^. Briefly, this method involves placing plastic tape under four electrodes (10-10 positions F5, F6, T7 and T8) to insulate these electrodes from the scalp, then adding an external wire to complete the circuit between the channel and the reference. Consequently, the activity recorded on these channels is the product of changes in magnetic flux. The EEG sampling rate was set to 5 kHz. Impedances were kept below 20 kΩ. All electrode positions, together with the nasion and left and right pre-auricular areas were digitised using a Polhemus Fasttrack system (Polhemus, Colchester, VT) for use in the creation of headmodels for source localisation.

### EEG preprocessing

All EEG analysis was carried out using MATLAB (MathWorks, Natwick, MA), the Fieldtrip^39^ and *fmrib*^40,41^ toolboxes, and custom scripts. The raw data was first high-pass filtered (1Hz; FIR). Following this, the gradient artifact was corrected using the FASTR algorithm implemented in the *fmrib* toolbox^40,41^. The gradient template for each TR was modelled on the average gradient artifact of the 60 nearest TRs. The data was then down-sampled to 500Hz and the ballistocardiogram (BCG) artifact was corrected using optimal basis set, again implemented in the *fmrib* toolbox. Heartbeat onsets were taken from the MR scanner’s physiological recordings. The continuous data was then inspected for large periods of movement which were marked and the associated MR scanner triggers deleted. Subsequently, the gradient and BCG corrections were repeated on the continuous data with the periods of movement excluded. This helped improve the accuracy of the gradient and BCG templates that were subtracted from the data. After gradient and pulse artefact correction the data from the motion sensors were used in a multi-channel recursive least squares algorithm to regress out the remaining movement-related artifacts^42,43^ (while retaining brain signal^44^) using custom scripts previously implemented by Jorge and colleagues^38^.

All subsequent EEG pre-processing was conducted using the Fieldtrip toolbox ^39^. First, the data was epoched into trials beginning 2 seconds before the onset of the video at perception/cue at retrieval and ending 4 seconds after the onset of the cue. Second, independent component analysis was used to remove blinks, saccades and any residual spatially-stationary noise that appeared to be linked to the cardiac artifact. Third, the data was demeaned, low-pass filtered (100Hz; Butterworth IIR) and re-referenced to the average of all channels. Fourth, the data was visually inspected to identify and reject any trials and/or channels containing residual artifacts (mean percentage of trials rejected: 23.1%; range: 10.4% to 39.1%). Fifth, the data was demeaned and re-referenced again to the average of all good channels (note that as any noise introduced by noisy channels in the earlier step will be shared by all good channels and therefore subtracted out during this re-referencing). Lastly, the scalp level data was reconstructed in source space to attenuate residual muscle artifacts (for details, see below).

### EEG source analysis

The preprocessed data was reconstructed in source space using individual head models, structural (T1-weighted) MRI scans and 4-layer boundary element models (BEM; using the dipoli method implemented in Fieldtrip). Electrode positions (as digitised via the Polhemus Fasttrack system) were mapped onto the surface of the scalp using fiducial points for reference. The timelocked EEG data was reconstructed using a Linearly Constrained Minimum Variance (LCMV) beamformer^45^. The lambda regularisation parameter was set to 5%.

### EEG time-frequency analysis

First, the source-reconstructed EEG data was convolved with a 6-cycle wavelet (−1 to 3 seconds, in steps of 25ms; 8 to 30Hz; in steps of 0.5Hz). Second, the resulting data was z-transformed using the mean and standard deviation of power across time and trials^8^. Third, the data was restricted to two time/frequency windows of interest (−1000-0ms and 500-1500ms post-stimulus; both 8-30Hz^23^) and then averaged across these windows, resulting in two alpha/beta power values per trial for each virtual electrode. To probe whether alpha/beta power decreased following stimulus onset these two values were contrasted in a one-tailed, non-parametric, cluster-based permutation-based t-test^46^ with 2000 randomisations. To investigate whether alpha/beta power decreased for remembered relative to forgotten trials, the data for the post-stimulus window was split by condition and contrasted using the same statistical approach.

### Combined EEG-fMRI analysis

An adjusted CVM approach outlined in *fMRI representational similarity analysis* was used to quantify information for this analysis. Rather than use a searchlight, CVM distance was computed in a region of interest (ROI) defined by the searchlight analysis. Specifically, this ROI consisted of all voxels included in any significant cluster revealed in the earlier analysis plus all neighbouring voxels that would have been included in the searchlight that contributed to the cluster. This approach maximised signal-to-noise for the measure of stimulus-specific information by only focusing on voxels where stimulus-specific information could be detected (see below for a note on circularity). As before, a training dataset was used to find weights that maximally discriminates two stimuli (per trial for encoding; averaged across repetitions for retrieval). In the case of retrieval data however, rather than project these weights onto stimulus-averaged testing dataset, these weights were projected onto the trial-level dataset. This change in approach provides a measure of stimulus-specific information for every trial within the specified ROI.

Similarly, an adjusted approach was used to quantify EEG power per trial. Whereas the prior section measured EEG power across all virtual electrodes, this analysis was restricted to virtual electrodes included in regions that coded for stimulus-specific information (as determined by the fMRI searchlight analysis). This approach ensured that the analysed EEG signal originated from the same region as the fMRI similarity index. Moreover, a ‘task-evoked’ decrease in alpha/beta power was calculated for each trial by subtracting the pre-stimulus alpha/beta power (−1000 to 0ms) from the post-stimulus alpha/beta power (500-1500ms). This reduced the influence of an increase in global alpha/beta power that typically occurs during the course of an experiment^47^.

These approaches yield a single measure of fMRI-derived stimulus-specific information and EEG-derived alpha/beta power for every trial. These fMRI and EEG measures were correlated across trials (separately for hits and misses in the case of the retrieval task). Given that the nature of the data of both variables is ratio, Pearson’s correlation was used. This returned an r-value for every participant, which underwent Fisher Z transform to approximate a normal distribution. Group-level statistical analysis saw these z-values being contrasted against the null hypothesis (z = 0; there is no correlation) in a one-tailed, non-parametric, permutation-based t-test^46^ with 2000 randomisations where the observed data and null hypothesis were permuted (again, separately for hits and misses in the case of the retrieval task). To test the spatial specificity of the effect, the correlation analyses above were re-run for each virtual electrode and then subjected to a one-tailed, non-parametric, cluster-based permutation-based t-test^46^.

We also addressed the spectral specificity of the effect. However, one should note that these results are difficult to interpret as both theta (3-7Hz) and gamma (40-50Hz) bands are much more susceptible to distortion by the MRI scanner than the alpha/beta band^26^. Aside from changes to the frequencies of interest, the analysis matched that which is described above. We considered both tails of the t-test, testing two differing hypotheses: 1) a reduction in power reflects an increase in information (mirroring the central hypothesis of the paper), and 2) as theta/gamma power typically increases during cognitive engagement^e.g.31^, an increase in power reflects an increase in information. This effect did not generalise to the theta (perception p = 0.228, retrieval p > 0.5) or gamma bands (perception p = 0.087, retrieval p = 0.097). To address the confound of confidence, a partial correlation was run on the recalled trials of retrieval task where confidence rating was included as a variable of no interest. The same partial correlation was not run on the perception data as we did not obtain a measure of confidence following the presentation of the video clip at perception. Similarly, a partial correlation was not run on the forgotten trials of the retrieval task as there was very little variation in the confidence measure (the majority of confidence ratings for forgotten trials equalled 1 [i.e. guess]) and hence would not prove to be a meaningful measure. To address the confound of BOLD activation, a partial correlation was run on all variants of the EEG * RSA correlation. The variable of no interest was BOLD activation of each trial, calculated by averaging the BOLD signal across voxels within the specified region of interest.

### A note on circularity

The use of data-driven regions of interest (ROIs) can, in some cases, introduce circularity into the analysis^48^. As a result, this can overestimate the size of an effect. However, we contend that our use of data-driven ROIs does not fall foul to this analytical flaw. Explicitly stated, the concern here is that by selecting the ROI that carries stimulus-specific information in the BOLD signal, we inflate the chance of finding a correlation between BOLD-derived stimulus-specific information and alpha/beta power in the same ROI. This concern is only valid when alpha/beta power also carries stimulus-specific information. In such an instance, we would essentially be limiting our correlation between two metrics of stimulus-specific information to a ROI where we know that (in this dataset) stimulus-specific information is represented. However, a Bayesian inference of RSA conducted on alpha/beta power (see results and section below) demonstrated that there is moderate evidence in favour of the null hypothesis that alpha/beta power does not carry stimulus-specific information. In light of this, we can infer that the use of data-driven ROIs in this instance does not introduce circularity into our analysis.

### EEG representational similarity analysis

To identify whether alpha/beta power carried stimulus-specific information, representational similarity analysis was conducted on the EEG time-frequency data (for perception and successful retrieval separately). The time-frequency data was derived in the same manner as described in the earlier section, but rather than average over time/frequency (as described in the third step), the individual time and frequency bins were retained. Representational similarity was quantified using Spearman’s correlation across all features (i.e. time, frequency and location) of every pair of trials. The resulting value underwent Fisher-z transformation to approximate a normal distribution. The observed similarity was then contrasted against the same models used in the earlier RSA approaches. This resulted in a single value describing stimulus-specific information for each subject, which was tested against the null hypothesis (there is no stimulus-specific information in alpha/beta power) in a one-tailed, non-parametric, permutation-based t-test^46^.

As we found insignificant evidence to support the alternative hypothesis, we then took a Bayesian approach to the statistical analysis. The same values used in above were analysed in a Bayesian one-sample t-test (as implemented in JASP, version 0.9 ^49^). We interpreted the resulting Bayes factor in line with the rule of thumb^50^.

