## Supplementary Materials for "Alpha/beta power decreases track the fidelity of stimulus-specific information"

*Univariate fMRI analysis*

The BOLD correlates of visual perception and episodic memory retrieval are well-documented. Here, we attempt to replicate these results to demonstrate the validity of our fMRI data. First, we investigated the BOLD correlates of visual perception by contrasting visual perception events with auditory perception events. To this end, a GLM was created per participant where these events were modelled as stick functions that had been convolved with a canonical hemodynamic response function (HRF). In addition, a regressor modelling button presses, six movement regressors and eight regressors modelling each run were added to the GLM. The derived beta weights for visual perception and auditory perception were then contrasted. The resulting contrast image of each participant was statistically appraised in a one-sample t-test across participants. Using a cluster-forming threshold of $p_{uncorr} < 0.001$ and $k = 10$, three significant clusters were identified: one in the occipital lobe ($p_{FWE} < 0.001$, k = 975, MNI [x = 42, y = -70, z = 10], Cohen's d = 2.09), one in the left temporal pole ($p_{FWE} = 0.008$, k = 67, MNI [x = -48, y = 2, z = -10], Cohen's d = 1.06), and one in the right temporal pole ($p_{FWE} = 0.005$, k = 72, MNI [x = 48, y = 5, z = -14], Cohen's d = 1.11) [see supplementary figure 1]. These results conform to earlier reports that visual stimulation produces greater activation in the occipital lobe than auditory stimulation[e.g.1].

Second, we investigated the BOLD correlates of visual memory retrieval by contrasting successful visual memory retrieval with successful auditory memory retrieval. The statistical approach matched that described above. Two significant clusters were identified: one in the left fusiform gyrus ($p_{FWE} = 0.001$, k = 89, MNI [x = 21, y = -37, z = -14], Cohen's d = 1.05), and one in the right temporal pole ($p_{FWE} = 0.001$, k = 99, MNI [x = -30, y = -46, z = -6], Cohen's d = 1.33) [see supplementary figure 1]. These results conform to earlier reports that visual memory retrieval is associated with greater activation of the fusiform gyrus than auditory memory retrieval[1].

Third, we further investigated the BOLD correlates of visual memory retrieval by contrasting successful visual memory retrieval with unsuccessful visual memory retrieval. The statistical approach matched that described above. Two significant clusters were identified: one in the occipital lobe ($p_{FWE} < 0.001$, k = 1178, MNI [x = 12, y = -52, z = -14], Cohen's d = 1.21), and one spanning the limbic system, including the hippocampus ($p_{FWE} < 0.001$, k = 1447, MNI [x = -21, y = -16, z = 2], Cohen's d = 1.33) [see supplementary figure 1]. These results conform to earlier reports of memory reactivation within sensory-specific regions[1] in conjunction with activation in the domain-general recollection network[2].

*EEG-confidence correlation*

It is plausible to suggest that the more information one recalls about an associated pair, the more confident they are in selecting the correct video. Therefore, it could be argued that alpha/beta power decreases are a confidence signal. To address this potential confound, we took our measure of EEG alpha/beta power (as derived in the same manner as described in the main text) and correlated this with the confidence rating provided on each trial. The derived r-value underwent Fisher z-transformation to approximate a normal distribution. These Fisher z-values were contrasted against the null hypothesis (there is no correlation; z = 0) across participants in a one-sample t-test. We found a significant negative correlation (p = 0.033, Cohen's d = 0.48), where a reduction in alpha/beta power was accompanied by an increase in confidence rating. While these results indicate a link between alpha/beta power and confidence, this link does not explain the link between alpha/beta power and stimulus-specific information (as evidenced by the partial correlation reported in the main text).

*Combined univariate EEG-fMRI analysis*

Here, we investigated the overlap between BOLD activity and the retrieval-related power decreases, complimenting earlier work looking at the overlap between BOLD and encoding-related power decreases[3]. To this end, a GLM was created in the same manner as in the *Univariate fMRI analysis* section with one key exception: the binary stick functions used in the earlier GLM were replaced with parametric values dictated by alpha/beta power observed on that trial. These parametric values were calculated by convolving the source-reconstructed EEG data with a 6-cycle wavelet (-1 to 3 seconds, in steps of 25ms; 8 to 30Hz; in steps of 0.5Hz). The resulting data was z-transformed using the mean and standard deviation of power across time and trials (for each condition separately[3]). Then, the data was restricted to the time/frequency window of interest (500-1500ms post-stimulus, 8-30Hz) and then averaged across this window and across all electrodes to return a single value of alpha/beta power per trial. Trials that were removed during preprocessing due to artifact contamination were given the value 0 in the parametric regressor. The statistical approach matched that of the *Univariate fMRI analysis* section. We uncovered two significant clusters where there was a greater negative relationship between BOLD and alpha/beta power for successfully recalled trials relative to forgotten trials: one in the occipital lobe ($p_{FWE} < 0.001$, k = 5183, MNI [x = -6, y = -76, z = 14], Cohen's d = 1.94), and the other in the parietal lobe ($p_{FWE} < 0.001$, k = 139, MNI [x = 39, y = -40, z = 38], Cohen's d = 1.33) [see supplementary figure 2].

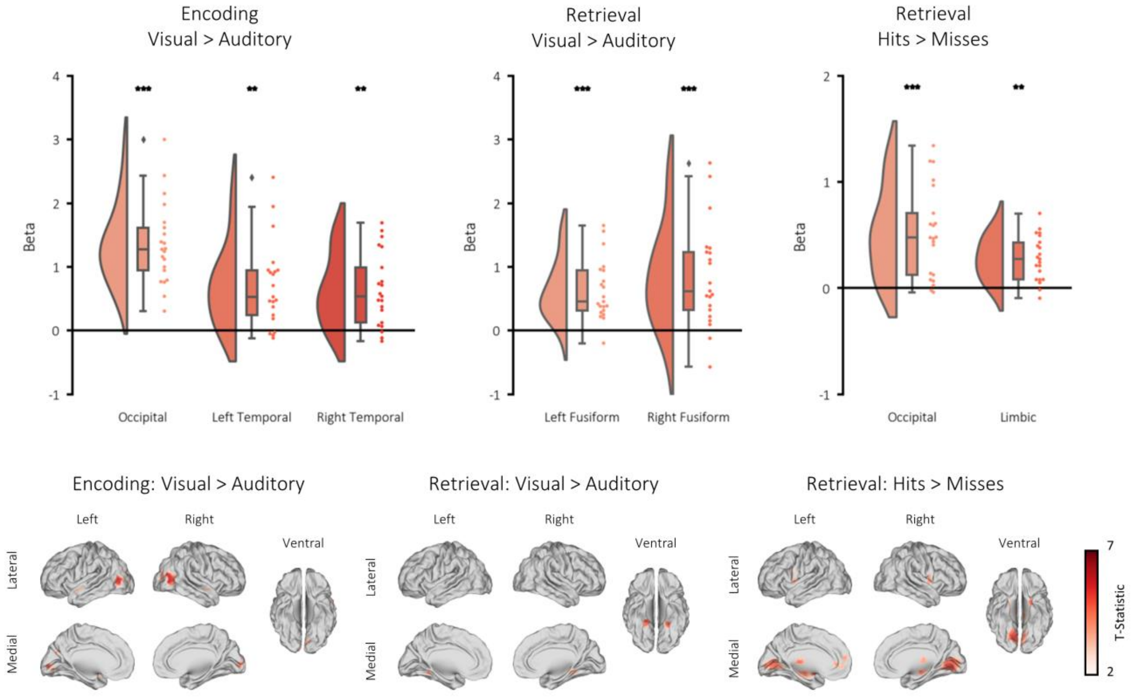


Supplementary figure 1. Univariate analysis of fMRI data. Top: raincloud plots depicting the contrasted beta weights for each participant for three tests: visual-specific perceptual activation (left), visual-specific retrieval-related activation (middle) and retrieval success (right). Bottom: brain maps depicting the clusters revealed from these contrasts (non-significant voxels masked).

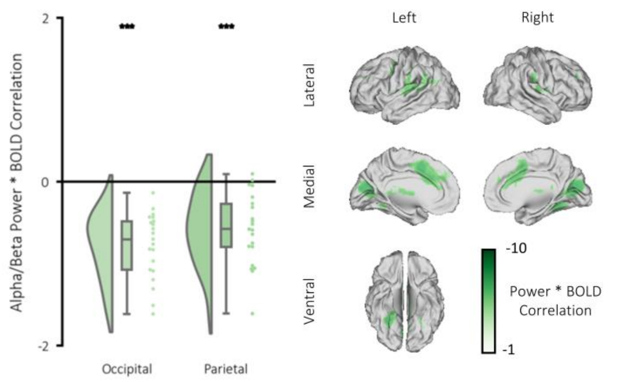


Supplementary figure 2. Correlation between alpha/beta power and BOLD signal. Left: raincloud plot depicting significant clusters identified when contrasting beta weights (remembered > forgotten) for each participant for three tests: visual-specific perceptual activation (left), visual-specific retrieval-related activation (middle) and retrieval success (right). Bottom: brain map depicting the clusters where alpha/beta power negatively correlated with BOLD for remembered items (relative to forgotten items).
